# Arginine methylation of the p30 C/EBPα oncoprotein regulates progenitor proliferation and myeloid differentiation

**DOI:** 10.1101/2024.03.28.587207

**Authors:** Linh T. Nguyen, Karin Zimmermann, Elisabeth Kowenz-Leutz, Dorothea Dörr, Anja Schütz, Jörg Schönheit, Alexander Mildner, Achim Leutz

**Affiliations:** Max-Delbrück Center for Molecular Medicine in the Helmholtz Association, Berlin, Robert-Rössle-Str. 10, 13125 Berlin, Germany; BSIO Berlin School of Integrative Oncology, Charité – Universitätsmedizin Berlin, corporate member of Freie Universität Berlin and Humboldt-Universität zu Berlin; Institute of Biomedicine at University of Turku, Turku, Finland; InFLAMES Research Flagship, University of Turku, 20014 Turku, Finland

**Keywords:** post-translational modification, transdifferentiation, myelopoiesis, leukemogenesis, GMP, AML

## Abstract

The transcription factor CCAAT enhancer binding protein alpha (C/EBPα) is a master regulator of myelopoiesis. *CEBPA* encodes a long (p42) and a truncated (p30) protein isoform from a single mRNA. Mutations that abnormally enhance expression of p30 are associated with acute myelogenous leukemia (AML). We show by mutational analysis that three highly conserved arginine residues (R140,147,154) located at the p30 C/EBPα N-terminus, previously found to be methylated, are involved in myeloid lineage commitment, progenitor proliferation, and differentiation. Replacement with lysine that retains the amino acid side chain charge enhanced progenitor proliferation, while uncharged side chains (alanine or leucine) impaired proliferation and enhanced granulopoietic differentiation. Analysis of protein-protein interactions (PPI) suggested that arginine methylation of p30 C/EBPα differentially determines its capacity to interact with SWI/SNF and MLL complexes. Pharmacological targeting of p30 C/EBPα arginine methylation may have clinical relevance in myeloproliferative and inflammatory diseases, in neutropenia, and in leukemic stem cells.

## INTRODUCTION

The C/EBPα transcription factor (TF) is an essential regulator of hematopoietic stem cell and progenitor biology. C/EBPα regulates myeloid lineage commitment, establishes the innate immune system from hematopoietic stem cells and multipotential progenitors (HSC, MPP) and is a crucial factor in myeloid leukemogenesis. Dysregulated expression and mutations of the *CEBPA* gene are associated with acute myelogenous leukemia (AML) and neutropenia (Heath *et al*, 2004; Pabst *et al*, 2001; Skokowa *et al*, 2009; Zhang *et al*, 2004).

*CEBPA* is a single-exon gene and produces a single mRNA, which encodes two divergent protein isoforms, the full length p42 and the truncated p30 C/EBPα. Translation of these variants can be initiated from alternative start sites in the same mRNA reading frame through the regulatory function of a small upstream open reading frame (uORF), located in the 5’ region of the mRNA (Calkhoven *et al*, 1994). Both isoforms induce myeloid lineage commitment of MPPs, yet p42 C/EBPα causes terminal cell differentiation and cell cycle arrest while p30 C/EBPα sustains progenitor proliferation (Nerlov, 2004).

Approximately 7-15% of AML patients carry mutations in the *CEBPA* gene that enable expression of p30 C/EBPα but abrogate expression or function of wild-type (WT) full length p42 C/EBPα. The function of dysregulated p30 C/EBPα as a driver oncoprotein was experimentally confirmed by targeted mouse genetics (Kirstetter *et al*, 2008). Mice and early progenitor cells that are deficient for both, p42 and p30 C/EBPα, lack granulocyte macrophage progenitors (GMP) and neutrophils, and are resistant to experimentally induced AML (Hayashi *et al*, 2011; Wesolowski *et al*, 2021). Expression of only the truncated p30 C/EBPα isoform from the murine *Cebpa* gene locus rescues GMP commitment and causes expansion of a myeloid progenitor population with complete leukemogenic penetrance (Bereshchenko *et al*, 2009). Thus, the C/EBPα p42, p30 isoforms display overlapping and antagonistic functions balancing myelopoiesis but the precise mechanism by which p30 encompasses both, lineage commitment and leukemogenic conversion remains elusive.

Both, p42 and p30 C/EBPα isoforms contain identical C-terminal basic DNA binding and leucine-zipper dimerization domains (bZip) and bind to the same regions in genomic DNA (Jakobsen *et al*, 2019). What distinguishes the C/EBPα isoforms is that p30 lacks parts of the p42 N-terminal tripartite transactivation domain (TAD, consisting of transregulatory elements TE1,2,3) and retains only TE3. Both C/EBPα isoforms interact with a plethora of proteins and protein complexes, however surprisingly, their interactomes are not only quantitatively different, but are also qualitatively distinguished, suggesting discrete regulatory capacity (Ramberger *et al*, 2021; Schmidt Heyes & Grebien, 2020).

The N-terminal regions of all CEBP-family members feature intrinsically disordered regions (IDR), comprising assemblies of conserved short linear motifs (SLiM) and molecular recognition elements (MoREs) that serve as multivalent combinatorial protein-protein interaction (PPI) modules to define their interactomes. The PPI modules and the flexible linker regions contain a plethora of post-translational modifications (PTM) that kaleidoscopically fine tune the interactomes and determine the resulting C/EBP biology (Dittmar *et al*, 2019; Leutz *et al*, 2011; Ramberger *et al*., 2021). In C/EBPα e.g., acetylation of specific lysine residues, such as K298 and K302, affect granulopoiesis while methylation of R35 directs the velocity of chromatin re-modelling and the acquisition of alternative myeloid cell fates (Bararia *et al*, 2016; Torcal Garcia *et al*, 2023).

Previously, we showed that the side chains of three highly conserved arginine residues in the N-terminus of p30 C/EBPα (human R142, 149, 156; here, rat R140, 147, 154) are targets of methylation and affect the interactome (Ramberger *et al*., 2021). Here, we used mutational analysis to examine whether amino acid side chain exchange affects p30 C/EBPα biology. Our data suggest that post-translational modifications of arginine side chains in p30 C/EBPα critically regulate its dichotomous activity in lineage commitment and progenitor expansion and may thus suite as future therapeutic targets.

## RESULTS

### Transdifferentiation capacity of p30 C/EBPα

The truncated isoform p30 C/EBPα is implicated in myeloid cell lineage commitment, early GMP expansion, and leukemogenicity (Kirstetter *et al*., 2008; Nerlov, 2004; Schmidt Heyes & Grebien, 2020). **Figure 1A** depicts an overview of the C/EBPα isoforms, the critical arginine residues R140,147,154 and the retroviral gene transfer constructs used in the experiments. A lymphoid-myeloid transdifferentiation (LMT) system was employed as a versatile tool to examine the relationship of C/EBP structure and function. The LMT system is based on v-Abl transformed mouse pre-B cells that are deficient for *Cebpa* and *Cebpb* genes (dKO-B, also termed B cells in the text), previously shown to prevent experimental complications due to cross regulation with endogenous C/EBP genes (Cirovic *et al*, 2017; Nguyen *et al*, 2024). We infected dKO-B cells with retroviral pMSCV-based EGFP-tagged p42 or p30 C/EBPα constructs. Cells were then examined by cytofluorometric analysis of CD11b surface antigen expression as an indicator of myeloid transdifferentiation (see below) and loss of the B cell marker CD19 (data not shown) to monitor conversion into myeloid cells, as schematically depicted in **Figure 1B, S1A**.

**Figure 1.**
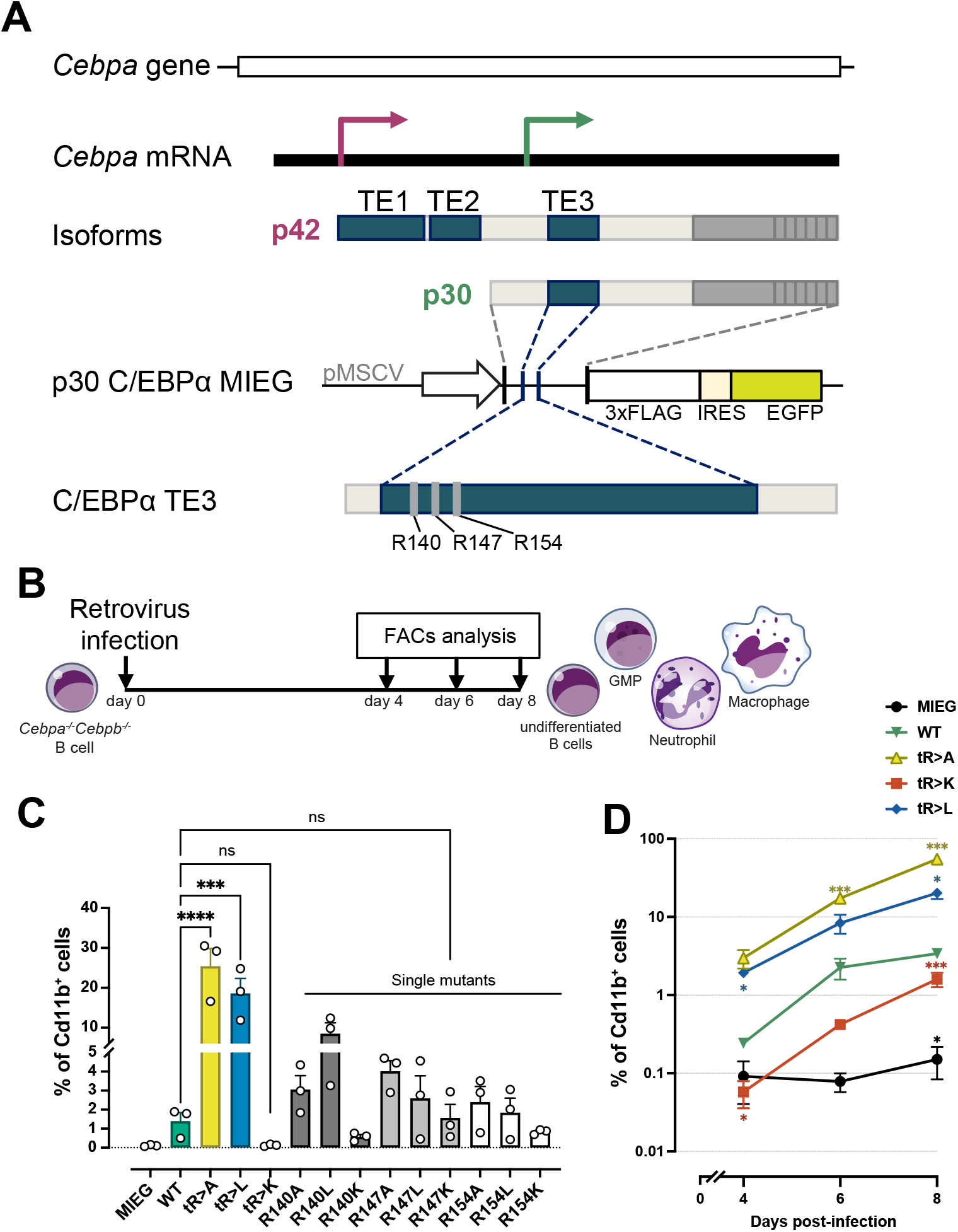
Lymphoid-myeloid transdifferentiation by WT and mutant p30 C/EBPα. **A.** Schematic representation of the C/EBPα gene, mRNA (top), protein isoforms and p30 C/EBPα expression constructs used in this study. The empty vector pMSCV-IRES-EGFP (MIEG) was used as control. Three arginine residues (R) of interest are shown. **B.** Schematic representation of the LMT assay. *Cebpa^-/-^Cebpb^-/-^*v-Abl transformed B cells (dKO-B cells) were retrovirally infected with constructs of interest. Aliquots of cell pools were harvested at indicated time points and analyzed by flow cytometry. **C.** Percentage of transdifferentiated myeloid CD11b^+^ cells induced by single and triple mutants after 6 days post-infection (pi). Data are shown as mean ± SEM, significance was determined by one-way ANOVA analysis followed by Dunnett’s multiple comparisons test, n=3, significance between WT p30 C/EBPα and mutants are shown. *p ≤ 0.05, **p ≤ 0.01, ***p ≤ 0.005, ****p ≤ 0.001. **D.** Percentage of CD11b^+^ cells at various time-points. Data are shown as mean ± SEM, significance was determined by two-way ANOVA analysis followed by Dunnett’s multiple comparisons test, n=4. Significance of comparisons to WT p30 C/EBPα and mutants are shown by asterisks in matching color, *p ≤ 0.05, **p ≤ 0.01, ***p ≤ 0.005, ****p ≤ 0.001.

As shown in **Figure 1C, S1B**, LMT conversion occurred quickly and profoundly not only after expression of p42 C/EBPα, but also slowly and sparsely with p30 C/EBPα, but not in controls. Cells that were transdifferentiated by p42 C/EBPα disappeared after 4-8 days due to arrest of proliferation, whereas p30 C/EBPα transdifferentiated myeloid cells persisted and accumulated over time (**Figure S1B, S1C**). These data confirm the capacity of p30 C/EBPα to induce myeloid lineage commitment without showing the growth inhibitory effect of p42 C/EBPα.

Next, we explored whether the three consecutive N-terminal arginine residues in p30 C/EBPα, previously found to be methylated, were involved in transdifferentiation and proliferation control. The arginine residues R140, R147, R154 of p30 C/EBPα were replaced individually or in combination by Alanine (A), Leucine (L) or Lysine (K). The effect on transdifferentiation of WT and mutant p30 C/EBPα was then compared, as shown in **Figure 1C**. In comparison to WT p30 C/EBPα, the triple mutants, tR>A and tR>L p30 C/EBPα, strongly enhanced the conversion into myeloid CD11b^+^ cells, while transdifferentiation by the triple tR>K mutant was impaired. The single site p30 C/EBPα mutants revealed that individual arginine side chain exchanges already, yet incrementally, altered transdifferentiation, in line with the stronger effects of the triple mutants. Although transdifferentiation by tR>K p30 was strongly impaired, prolonged cultivation of R3K p30 C/EBPα infected cells nevertheless stably maintained a small population of CD11b^+^ myeloid cells, as shown in **Figure 1D**.

The data suggested that the lineage commitment and differentiation capacity of p30 C/EBPα is gradually regulated by modifications of three N-terminal arginine residues. Overall, the triple mutants strongly augmented the phenotypes, as seen by the individual amino acid replacements, suggesting that the three respective side chains function in co-operation.

### Transcriptomic profiling of p30 C/EBPα transdifferentiated cells

To further delineate the cellular characteristics of transdifferentiated cells generated by WT and mutant variants of p30 C/EBPα, we isolated GFP^+^ cells at day 4 post-infection and subjected them to bulk RNA sequencing analysis (**Figure 2A**). Principal component analysis (PCA) demonstrated distinct differences between WT, the p30 C/EBPα triple mutant variants, and the control group (**Figure 2B**), indicating that all p30 C/EBPα variants entail the capability to induce distinct cellular identities. In comparison to B cell controls, the cells expressing the tR>A and tR>L p30 C/EBPα exhibited the most pronounced transcriptomic alterations, while the WT p30 C/EBPα occupied an intermediate state, between tR>A, tR>L on the one hand and tR>K p30 C/EBPα expressing cells, on the other (**Figure 2B**).

**Figure 2.**
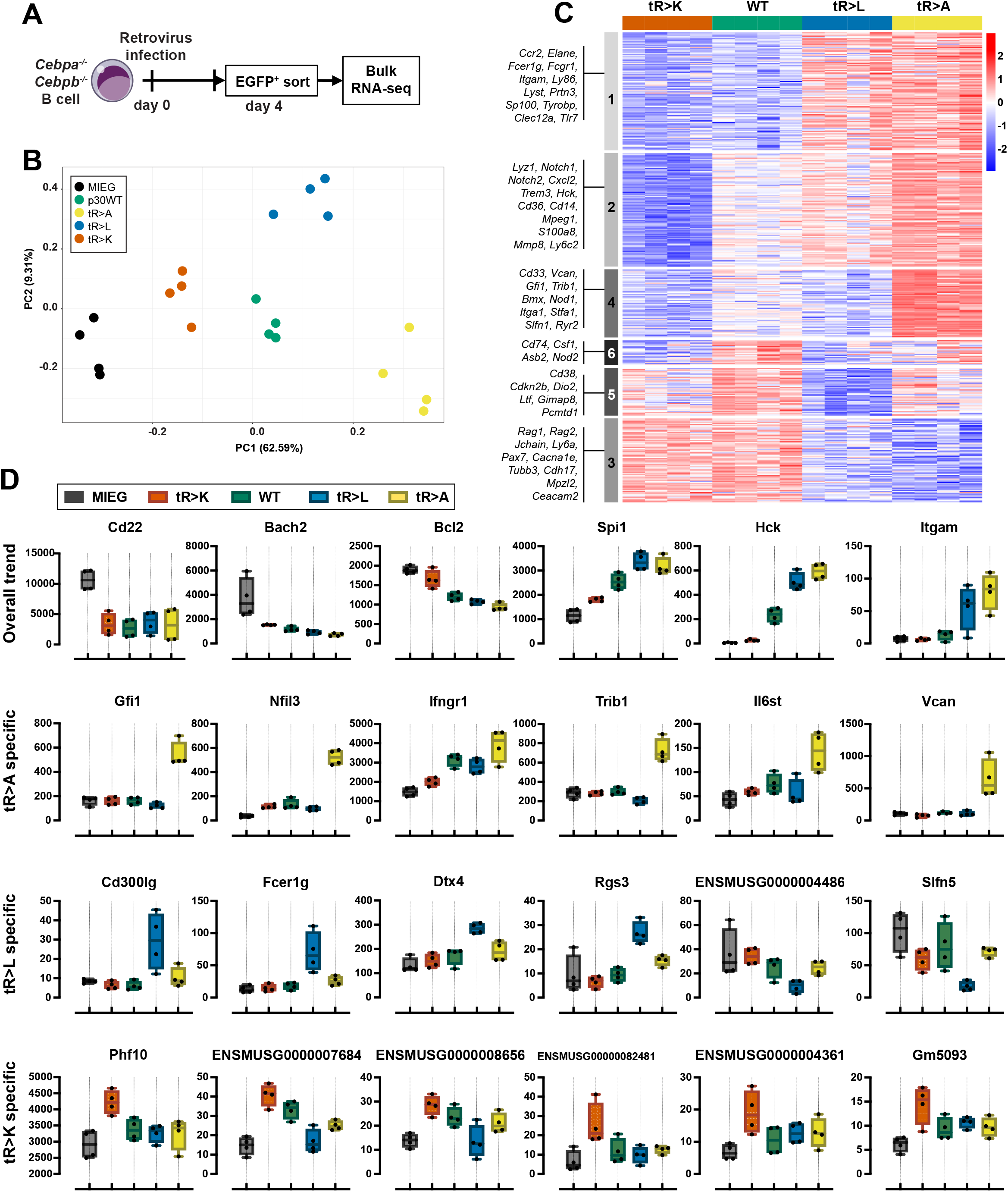
Transcriptomic profiling of WT and mutant p30 C/EBPα cells. **A.** Schematic representation of sample preparation for bulk RNA-seq. **B.** Principal component analysis of the top 500 differentially expressed genes. Constructs are color coded and shown in inset on the left. **C.** Heatmap presenting differential gene expression by constructs as indicated on the top. Color code as in B. Genes with adjusted p-value 0.05, |FC|>2 in at least one comparison are shown. Representative gene names of each cluster are listed on the left. **D.** Expression of representative genes, which were manually selected based on read counts. Top row: genes expressed in overall trends, last 3 rows: genes expressed in mutant-specific patterns.

Differential myeloid transdifferentiation process became evident by the enrichment of gene ontology (GO) terms associated with myelomonocytic features, such as chemotaxis and the production of Interleukin (IL)-1 or IL-6. Notably, residual transdifferentiation capacity was observed in the tR>K p30 C/EBPα variant, as evidenced by the GO term "Monocyte chemotaxis" (**Figure S2A**).

Pairwise examination of differential gene expression identified 1273 genes that exhibited significant alterations (adjusted p-value 0.05, |FC|>2) in at least one of the comparisons (**Figure S2B**). The global gene expression profile was consistent with the findings obtained by flow cytometric analysis of the myeloid marker CD11b (**Figure 1C**). Specifically, B cell-associated genes (*Cd22, Bach2, Bcl2*) exhibited high expression levels in vector control cells, slightly reduced expression trend in p30 tR>K-expressing cells, followed by p30 WT-, tR>L- and strong reduction in tR>A-expressing cells (**Figure S2B; 2C, cluster 3**). Conversely, the expression of characteristic myeloid genes, encompassing direct C/EBPα target genes, transcription factors, surface markers, and cytokines, exhibited an inverse correlation. Myeloid gene expression was highest in cells expressing the tR>A and tR>L p30 C/EBPα mutants, less pronounced in p30 WT-expressing cells, and lowest in tR>K p30 C/EBPα expressing cells (**Figure S2B; 2C, cluster 1, 2**).

Comparisons between the WT p30 C/EBPα and each of the triple mutant variants revealed noteworthy distinctions, as shown in the cluster analysis in **Figure 2C**. While all three, namely WT, tR>A, and tR>L p30 C/EBPα mutants, demonstrated the capacity to induce myeloid lineage switching, the tR>A and tR>L expressing cells exhibited a higher abundance of genes associated with granulocyte/neutrophils (*Elane, Prtn3, S100a8, Mmp8, Gfi1*), as compared to WT p30 C/EBPα (**Figure 2C**, particularly clusters 1, 2, 4). On the other hand, clusters 5 and 6 represent sets of genes that were either marginally or not induced by the tR>L p30 C/EBPα mutant (**Figure 2C, cluster 5, 6)**. These included genes that are specific to monocytes (cluster 6) and a collection of mitochondrial pseudogenes of unknown function (cluster 5). In contrast, both the tR>K and WT p30 C/EBPα variants exhibited a higher expression of genes specific for B cells (*Rag1/2, Jchain, Ly6a*), depicting the attenuated capacity of tR>K p30 C/EBPα to induce myeloid lineage switching.

Next, we searched for mutant specific differentially expressed genes that that play a role in myelopoiesis but deviated from the overall trends. The tR>A and tR>L p30 C/EBPα mutants exhibited upregulation of the known C/EBPα target genes, *Trib1* and *Dtx4*, respectively. Furthermore, they augmented expression of myeloid transcripts, including *Gfi1*, *Flt3*, and *Vcam,* in the case of tR>A, and *CD300lg*, *Fcer1g*, and *Rgs3* in the case of tR>L **(Figure 2D).** In contrast, the tR>K p30 C/EBPα mutant induced a small number of pseudogenes (**Figure 2D)**, which may potentially be linked to unknown regulatory mechanisms (Cheetham Faulkner & Dinger, 2020).

In summary, the data show that alterations of arginine side chain positions R140,147,154 have a significant impact on the transdifferentiation potential of p30 C/EBPα. In comparison to WT p30 C/EBPα, mutations that result in the loss of charge (R to A, L substitutions) or mimic methylation (R to L) promote myeloid transdifferentiation, whereas R to K mutations, which retain side chain charges but obliterate arginine specific methylation, constrained transdifferentiation.

### Phenotypic potential of p30 C/EBPα mutants in progenitor cells

In the absence of p42, p30 C/EBPα induces myeloid commitment and enhances leukemic expansion of progenitor cells, refelcting its functions as a regulator of cell fate and as a myeloid driver oncoprotein (Kirstetter et al., 2008a). We therefore examined the functional characteristics of WT and p30 C/EBPα mutant variants using serial colony replating of early progenitor cells derived from mouse bone marrow (Lineage depleted, Sca1 positive, c-Kit negative, LSK; bone marrow cells derived from Cebpa*^fl/fl^*Cebpb*^fl/fl^* “WT” mice) (Ikuta & Weissman, 1992; Li & Johnson, 1995). LSK cells were transduced with p30 C/EBPα or control vector constructs and GFP^+^ sorted cells were plated in semi-solid medium supplemented with cytokines and replated following a standard regime to determine how p30 constructs affect the replication capacity and colony characteristics (Eaves, 2015).

**Figure 3A** schematically shows the examination of the colony morphology, clonogenicity and proliferation capacity. Serial replating assays of c-Kit-enriched cells showed that GFP^+^ cells expressing tR>K p30 C/EBPα maintained replating capacity up to or beyond the fourth passage, whereas all other p30 C/EBPα constructs led to loss of clonogenicity after the second and third rounds of replating **(Figure S3A).** We next compared side-by-side the serial replating capacity of early progenitor LSK cells and GMP cells (defined as Lineage^-^Sca-1^-^c-Kit^+^FcgRIII^+^, **Figure S3B**) expressing p30 C/EBPα mutants. For both hematopoietic developmental stages, the tR>K p30 C/EBPα mutant exhibited similar replating capabilities up to the fourth passage (**Figure 3B, 3C**). However, the cells derived from the LSK displayed more colonies in comparison to GMPs, indicating the superior clonogenicity of very early progenitors.

**Figure 3.**
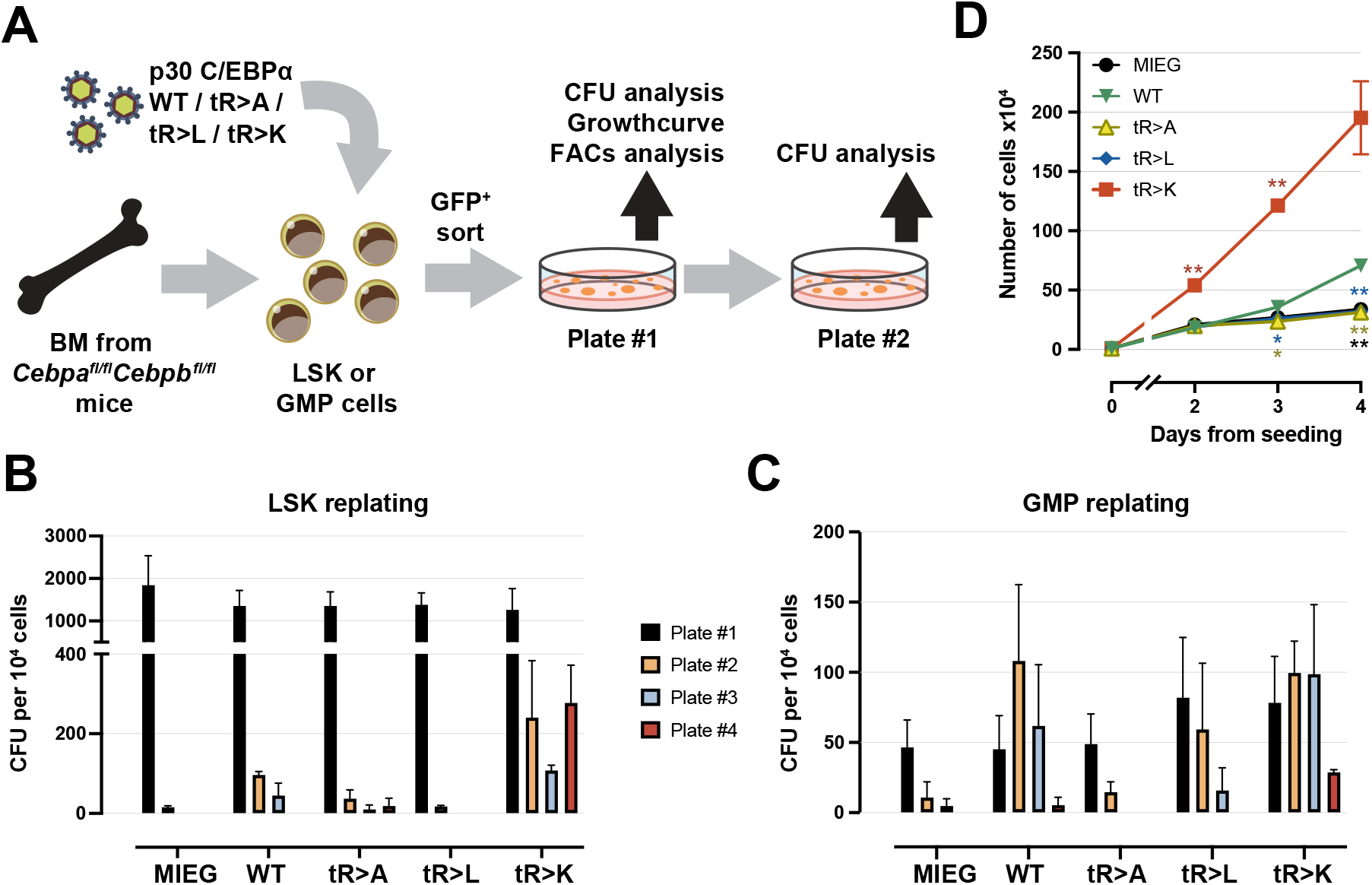
Serial replating potential of progenitor cells expressing WT or mutant p30 C/EBPα. **A.** Schematic representation of serial replating assay. Bone marrow cells were harvested from *Cebpa^fl/fl^ Cebpb^fl/fl^* mice. Cells were enriched for c-Kit using magnetic cell sorting, followed by LSK and GMP flow cytometry sorting. Retroviral infections were performed 24 hours after cell harvesting (day 0). GFP^+^ cells were sorted at day 2-pi and plated. Colonies were counted/re-seeded every 7 days. **B-C.** Colony formation recorded over four passages of replating LSK cells **(B.)** and GMP cells **(C.)**. Colony numbers were determined by manual counting on scanned microscopic images. Colony formation rates per 10^4^ cells from three biological replicates are shown. **D.** Growth curve of accumulative cell counts using cells derived from Plate #1 of total c-Kit enriched bone marrow replating (as shown in Figure S3A). Data are shown as mean ± SEM, significance was determined by two-way ANOVA analysis followed by Dunnett’s multiple comparisons test, as shown by asterisks in matching color, *p ≤ 0.05, **p ≤ 0.01, ***p ≤ 0.005, ****p ≤ 0.001. Results of one experiment with three technical replicates are shown. Three independent experiments were performed with similar results.

Cells derived from the starting c-Kit-enriched culture (Plate #1, **Figure 3A, S3A**) were further examined by proliferation analysis, including cell counting, colorimetric (WST-1) proliferation assay, and dye dilution assay. Notably, the tR>K p30 C/EBPα mutant expressing cells exhibited the highest increase in cell number (**Figure 3D**) and proliferation (**Figure S3C, S3D**) across all three assays, as compared to the other p30 C/EBPα variants. The tR>K p30 C/EBPα mutant also exhibit faster cytoplasmic dye dilution (CellTrace Violet, **Figure S3D**), indicating acceleration of the cell cycle. The WT p30 C/EBPα construct also induced proliferation and clonogenicity, although to a lesser extent than that of tR>K. Collectively, the results suggest that among the p30 C/EBPα constructs tested, the tR>K p30 C/EBPα variant augment both the clonogenic potential and the proliferation of progenitor cells at the HSC and GMP stages.

Next, we examined the fates of cells derived from LSK colonies (Plate #1, **Figure 3A, S3A**) that expressed WT or mutant p30 C/EBPα. Four morphological categories were used to determined colony types, as depicted in **Figure 4A**. The predominant type of colonies induced by p30 C/EBPα tR>L exhibited a typical granulocytic CFU-G morphology, characterized by a compact core surrounded by dispersed small cells (**Figure, 4B**). In contrast, most colonies generated by tR>K p30 C/EBPα resembled monocytic CFU-M, with highly dispersed colonies, containing large cells (**Figure 4B**).

**Figure 4.**
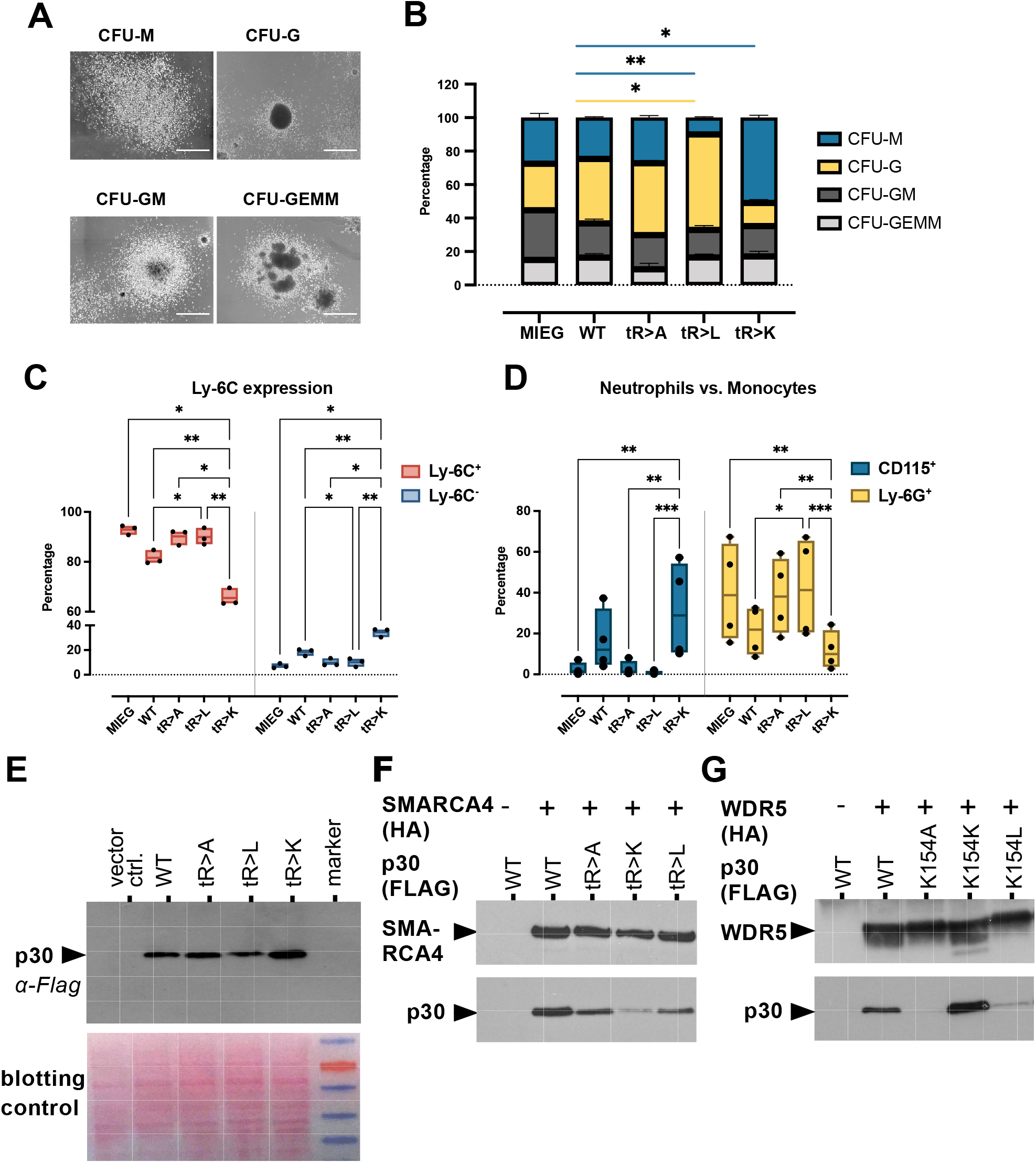
Myeloid differentiation induced by WT and mutant p30 C/EBPα. **A.** Characteristic types of colony morphologies on Plate #1. Scale bar 100 μm. Determination of colony types was as listed in Thermo Fisher Scientific (provider of methylcellulose based medium M3434 for colony formation assay used in this experiment). **B.** Percentage of colony types on Plate #1. Constructs are indicated below stacked bar graphs. Colonies were counted using scanned images and categorized according to **A.**. Representative results from one experiment with three technical replicates are shown. Data are shown as mean ± SEM, significance was determined by two-way ANOVA analysis followed by Dunnett’s multiple comparisons test. Only significant comparison between WT and other constructs is shown, *p ≤ 0.05, **p ≤ 0.01, ***p ≤ 0.005, ****p ≤ 0.001, color code representing colony types are shown on the right. **C-D.** Percentage of Ly6C^-^ and Ly6C^+^ population (**C.**) and Ly6C^+^CD115^+^ monocytes and Ly6C^+^Ly6G^+^ neutrophils (**D.**) on Plate #1. Data are shown as mean ± SEM, n=3, significance was determined by two-way ANOVA analysis followed by Dunnett’s multiple comparisons test, *p ≤ 0.05, **p ≤ 0.01, ***p ≤ 0.005, ****p ≤ 0.001, only significant comparisons are shown. **E.** Upper panel: Expression of WT and triple mutant p30 C/EBPα constructs in retroviral packaging Plat-E cells. Protein lysates were separated by SDS-gel electrophoresis, blotted and probed with anti-FLAG to reveal p30 C/EBPα expression, as indicated. Lower panel: Ponceau staining as lysate and blotting control of the same blot. **F.** Interaction of WT or triple mutant p30 C/EBPα with SMARCA4. HEK293T were co-transfected with SMARCA4-HA and C/EBPα-FLAG constructs, as indicated. SMARCA4 was immunoprecipitated and detected with anti-HA. Co-immunoprecipitated WT or triple mutant p30 C/EBPα were detected by anti-FLAG. **G.** Interaction of WT or single mutant p30 C/EBPα with WDR5. Co-immunoprecipitation protocol as in **F.**, from cells co-transfected with WDR5-HA and C/EBPα-FLAG constructs, as indicated.

Thus, the emergence of CFU-G and CFU-M type colonies correlated with the tR>L and tR>K versions of p30 C/EBPα, respectively. Cytology of cells, as revealed by cytospins obtained from pooled colony populations, further supports these observations, showing a high proportion of typical neutrophil granulocytes at different maturation stages (bi- or multilobed, polymorphic nuclei, cytoplasmic azurophilic granules) with tR>L p30 C/EBPα cells while the majority of tR>K p30 C/EBPα were monocytes (rounded shape, pale blue- gray cytoplasm, small lilac granules and vacuoles) (**Figure S4A**).

Cells from colony Plate #1 were also analyzed by flow cytometry to distinguish the distribution of granulocytes (Ly6C^+^Ly6G^+^CD115^-^) and classical monocytes (Ly6C^+^Ly6G^-^ CD115^+^) (**Figure S4B**). We noticed an accumulation of Ly6C^-^ cells in colonies with tR>K p30 C/EBPα, while these cells occupy less than 20% in the other groups (**Figure 4C**). Within Ly6C^+^ population, differentiation towards Ly6G^+^ granulocytes was pronounced with tR>L p30 C/EBPα, while CD115^+^ monocyte differentiation was favored with tR>K p30 C/EBPα (**Figure 4D**). The phenotypes induced by WT p30 C/EBPα fell between those induced by tR>A or tR>L p30 C/EBPα on one end of the spectrum and tR>K p30 C/EBPα on the other. Notably, the accumulated Ly6C^-^ population in tR>K p30 C/EBPα cells exhibited no overlap with other myeloid marker expressing populations (Ly6G, CD115, Cd11b, F4/80), ruling out the possibility they represent non-classical Ly6C^lo/neg^ monocytes, instead they likely represented an undifferentiated population (**Figure S4C**).

Finally, we examined whether the undifferentiated Ly6C^-^ population accounted for the enhanced proliferation of tR>K p30 C/EBPα expression cells using a dye dilution assay. Staining of Ly6C was integrated in the assay shown in **Figure S4D**. The dye dilution rate of Ly6C^+^ and Ly6C^-^ subsets showed minimal differences between tR>A, tR>L p30 C/EBPα expressing cells, while tR>K and WT p30 C/EBPα expressing cells showed higher division rate in the Ly6C^+^ subset (**Figure S4D**). In both Ly6C subsets, proliferation was highest with tR>K (**Figure S4D**). During a 7-day time course, we observed an overall trend in that the Ly6C^+^ population gradually diminished (in population percentage and cell fitness, microscopic observation, data not shown), while the Ly6C^-^ cells maintained growth. This implies that beside cell cycle progression, prevention of cell death may also play a role in tR>K p30 C/EBPα mutant cells. Altogether the data suggest that the tR>K p30 C/EBPα mutant supports proliferation of hematopoietic precursors and myeloid progenitors at various differentiation stages.

Seminal genetic studies conducted in Drosophila melanogaster and Saccharomyces cerevisiae have unveiled numerous components of the Trithorax/COMPASS-like/MLL and SWI/SNF/BAF complexes. These complexes, initially identified for their roles in segmental transformation and mating-type switching, have since been recognized as multi-component protein complexes involved in chromatin modification and remodeling, respectively. Both complexes are categorized within the Trithorax-group, a set of evolutionarily highly conserved epigenetic regulators crucial for cell type specification, including hematopoiesis and leukemogenesis (Cenik & Shilatifard, 2021; Rodrigues Shvedunova & Akhtar, 2020).

Previous studies have demonstrated physical and functional interactions between both complexes and C/EBPα during critical developmental processes in hematopoietic differentiation and proliferation (Grebien *et al*, 2015; Ohlsson *et al*, 2014; Wesolowski *et al*., 2021). In an endeavor to integrate our biological findings with previously reported interactions, we investigated the core components of SWI/SNF and MLL complexes for their interaction with wild-type (WT) and mutant p30 C/EBPα using co-immunoprecipitation assays, as depicted in **Figure 4E-G**.

The interaction between SWI/SNF and C/EBPα has been previously documented to rely on the N-terminus of p30 C/EBPα (Muller *et al*, 1999; Muller *et al*, 2004; Pedersen *et al*, 2001). Additionally, proteomic analyses have shown an enrichment of SWI/SNF with the tR>L p30 C/EBPα mutant and methylated R140 over the wild-type (WT), including the core ATPase subunit SMARCA4 (Brm) (Ramberger *et al*., 2021). As illustrated in **Figure 4E, F**, co-immunoprecipitation confirmed that both differentiation enhancing tR>L, tR>A p30 C/EBPα mutants exhibited a stronger interaction with SMARCA4 (Brm) compared to the proliferation-enhancing tR>K p30 C/EBPα mutant, which displayed a reduced interaction.

Progenitor proliferation induced by p30 C/EBPα relies on the recruitment of the MLL complex through the adapter protein WDR5, which is also a constituent of several other chromatin regulatory complexes (Grebien *et al*., 2015; Guarnaccia & Tansey, 2018; Ohlsson *et al*., 2014; Schmidt *et al*, 2019; Wesolowski *et al*., 2021; Wysocka *et al*, 2005). To investigate the interaction of WDR5 with differentially methylated p30 C/EBPα N-termini, we employed the motif-based screening method of immobilized tiled peptides (PRISMA; utilizing both WT and post-translational modification-modified tiled peptide configurations) (Dittmar *et al*., 2019). As depicted in **Figures 4G and S4E-H**, WDR5 exhibited high specificity in binding to peptides containing the unmethylated side chain configuration of R154 or to positive control peptides derived from histone H3 and MLL (Song & Kingston, 2008; Wysocka *et al*., 2005). Symmetric or asymmetric methylation at position R154 of p30 C/EBPα peptides, or substitution with alanine or citrulline, hindered the binding of WDR5, thus confirming the selectivity and methylation sensitivity of these interactions. Finally, co-immunoprecipitation assays validated the association of WDR5 with both WT and R154K p30 C/EBPα in cells, whereas the R154A and R154L p30 C/EBPα mutants exhibited reduced association with WDR5 (**Figure 4G**).

In summary, these findings highlight how distinct arginine residues and their modification status in p30 C/EBPα differentially influence the interaction with critical components of the MLL and SWI/SNF complexes.

## DISCUSSION

Contrary to the prevailing concept that p30 C/EBPα primarily functions as a dominant-negative variant of the full-length p42 C/EBPα, our data show that p30 C/EBPα instead can direct cell fate and progenitor proliferation in a PTM-dependent fashion. Three conserved arginine residues (R140, 147, 154) located in the N-terminus of p30 C/EBPα are critically involved and point mutations at these sites distinctly modify the biological functions of p30 C/EBPα in lineage commitment, differentiation, and proliferation. Our data add evidence to the concept that arginine side chain methylation of pioneering TF can determine epigenetic outcomes upstream of histone modifications (Leutz *et al*., 2011; Saber & Rudnicki, 2022; Torcal Garcia & Graf, 2021).

The most plausible explanation for our data is that the p30 C/EBPα mutants represent hypo- and hypermorphic mimics of arginine methylation. Accordingly, tR>K p30 C/EBPα represents the unmethylated state of WT p30 C/EBPα and facilitates progenitor expansion and monocytic cell fate, while methylation of arginine side chains, represented by tR>L p30 C/EBPα, is associated with restrained progenitor expansion and granulocyte differentiation. This interpretation aligns with previous biochemical data demonstrating that the arginine methylation status of C/EBPs profoundly alters their interactomes and biological functions (Dittmar *et al*., 2019; Kowenz-Leutz *et al*, 2010; Ramberger *et al*., 2021; Torcal Garcia *et al*., 2023). Hence, forthcoming studies decoding the upstream and downstream regulatory p30 C/EBPα network in hematopoietic progenitor biology and leukemia should consider the cellular machinery involved in C/EBPα arginine methylation.

The tR>A,L p30 C/EBPα mutant and the tR>K elicit opposing phenotypes. The tR>A,L mutants promote granulopoietic differentiation and attenuate clonogenicity and proliferation. In contrast, the tR>K p30 C/EBPα mutant, that retains the positively charged side chains, supports proliferation and elicits a monocytic phenotype. The WT p30 C/EBPα settles between both phenotypes, in consistence with its arginine side chains in either unmodified or (partially) modified configuration. In accordance, methylation of human R142 (equivalent to rat R140) and the tR>L mutant were previously shown to enhance interaction with SWI/SNF, whereas methylation of R154 inhibits binding of WDR5 that mediates binding of the MLL complex (Grebien *et al*., 2015; Ramberger *et al*., 2021; Schmidt *et al*., 2019). Thus, the differential interactions with SWI/SNF and MLL complies with the state of arginine modifications and with the biology of the p30 C/EBPα mutants, suggesting an important mechanistic connection.

Interestingly, the p30 C/EBPα triple mutants amplify the phenotypes that were already observed with single-site mutants (see **Figure 2**), suggesting individual and combinatorial functions of the adjacent arginine side chains. How can we reconcile the functional co-operativity of adjacent arginine PTMs with their individual specificity? PTM-dependent graduated non-stoichiometric functions are common biochemical features of linear regulatory protein motifs, including SLiMs (Holehouse & Kragelund, 2024; Moses Hériché & Durbin, 2007; van der Lee *et al*, 2014). In the case of C/EBPα, we envisage that differential arginine modifications may “titrate” the reorganization propensity e.g. by structural “breathing” of the p30 C/EBPα N-terminus to modulate the access and function of its multivalent TE3 - PPI motifs and, consequently, enable alternative biological outcomes. Considering the recent observation that C/EBPα undergoes biomolecular condensation that depend on aromatic amino acid in the intrinsically disordered N-terminal region, one may also consider phase-separation processes to be involved that are sensitive to arginine methylation (Christou-Kent *et al*, 2023). Notably, three highly conserved tyrosine residues (Y129/136/145) are located in the vicinity of the triple arginine residues in p30 C/EBPα. Alternating R-Y residues are a typical feature of IDRs that engage in non-covalent pi-cation side chain interactions during polypeptide coalescence that are disrupted by arginine side chain methylation. Although speculative, this notion may explain why both, leucine and alanine substitutions of R140/147/154 p30 C/EBPα display similar biological outcomes, as neither alanine nor leucine side chains engage in pi-cation based interactions, while replacement with lysine can retain this function (Crowley & Golovin, 2005; Fisher & Elbaum-Garfinkle, 2020; Gallivan & Dougherty, 1999; Wang *et al*, 2022).

### Conclusion and Limitations of the Study

The findings presented in this study provide new insights into key functions of the truncated p30 C/EBPα isoform and may hold promise for future targeted pharmacological interventions in p30 C/EBPα driven leukemia. However, the observations also raise questions about the dynamics and the enzymatic machinery involved in p30 C/EBPα methylation, its connections to cytokines and signaling pathways, intracellular transmission cascades, the underlying molecular genetics of chromatin structure, gene regulation, and homeostatic or leukemic effects in the living organism. As our studies are currently limited to in vitro assays, important future tasks will have to address C/EBP modifications in relation to normal hematopoiesis and disease in the living organism.

Furthermore, investigating the impact of the respective p30 C/EBPα arginine PTMs in the p42 C/EBPα context will be important. Further, it will also be important to consider the positional isomeric status and the degree of arginine modifications, i.e. mono-, symmetric, and asymmetric arginine di-methylation, in relation to upstream signals and downstream effects, in analogy to what has been termed “histone code” in chromatin regulation.

Moreover, methylation of arginine side chains appears to be largely irreversible, suggesting it as an regulatory end point, except that citrullination by peptidyl arginine deiminases (PADI) may remove methylated and unmethylated arginine side chains to create a new side chain configuration (Tilvawala & Thompson, 2019).

In summary, the results shown here serve as a foundation and starting point, more comprehensive research on the connections between p30 C/EBPα PTM in the intact organism and its biological impact will be required. While the current study falls short of a holistic analysis, our data provide a new rational of the dichotomous function of p30 C/EBPα in progenitor expansion and differentiation. This study may encourage further investigations of the physiological placement and impact of p30 C/EBPα PTM and its turnover in normal hematopoiesis and disease and as a pharmacological target.

## MATERIALS AND METHODS

### Resource availability

#### Materials availability

*Corresponding authors:* Achim Leutz

Requests for materials, additional raw and analyzed data will be reviewed by the corresponding author to verify if the request is subject to any intellectual property of confidentiality obligations. Any data and materials that can be shared will be released via a material transfer agreement.

#### Data and code availability

The RNA sequencing data are publicly available at the National Center for Biotechnology Information with Gene Expression Omnibus accession no.

### Cell culture

The LMT system employed the Cebpa^Δ/Δ^Cebpb^Δ/Δ^ v-Abl transformed pre-B cell line generated as described before (Cirovic *et al*, 2017). The established clone 18 of this cell line were used throughout this study and termed dKO-B cells. dKO-B cells were maintained in complete DMEM (10% fetal bovine serum (FBS), 10 mM HEPES and penicillin/streptomycin) supplemented with 50 mM β-mercaptoethanol. For retrovirus generation, Plat-E was used as packaging cell line. Bone marrow-derived progenitors and stem cells were culture in IMDM supplemented with 10 ng/mL IL-3, 10 ng/mL IL-6 and 20 ng/mL mSCF.

### Expression constructs and retroviral infection

C/EBP genes were expressed using pMSCV-IRES-EGFP (MIEG) retroviral vectors as described (Cirovic *et al*, 2017). Point mutations in *Cebpa p42* were introduced by site-directed mutagenesis using the QuikChange Site-Directed Mutagenesis Kit (Stratagene #200518). Primers used for single mutant exchange were: R140A forward 5’- ggctacctggacggcgcgctggagcccctgta and reverse 5’-tacaggggctccagcgcgccgtccaggtagcc; R140L forward 5’-ggctacctggacggcctgctggagcccctgta and reverse 5’- tacaggggctccagcaggccgtccaggtagcc; R140K forward 5’- ggctacctggacggcaagctggagcccctgta; R147A forward 5’-gagcccctgtacgaggccgtcggggcgcccgc and reverse 5’-gcgggcgccccgacggcctcgtacaggggctc; R147L forward 5’-gagcccctgtacgacctcgtcggggcgcccgc and reverse 5’- gcgggcgccccgacgaggtcgtacaggggctc; R147K forward 5’- gagcccctgtacgacaaagtcggggcgcccgc; R154A forward 5’- ggggcgcccgcgctggcgccgctggtgatcaa and reverse 5’-ttgatcaccagcggcgccagcgcgggcgcccc; R154L froward 5’-ggggcgcccgcgctgctgccgctggtgatcaa and reverse 5’- ttgatcaccagcggcagcagcgcgggcgcggg; R154K forward 5’- ggggcgcccgcgctgaagccgctggtgatcaa and reverse 5’-ttgatcaccagcggcttcagcgcgggcgcggg. Mutated C/EBPa p42 constructs were used as templates for synthesizing p30 C/EBPα constructs using polymerase chain reactions and EcoRI/XhoI restriction site-targeted primers: forward 5’-gcgaagcttgaattcgccatggcggcgggggcgcacggc and reverse 5’- ccgctcgagctagagcttgtcatcgtcatccttgtaatc. p30 C/EBPα fragments were cloned into MIEG to generate pMSCV_p30-mutant_3xFLAG_IRES_EGFP vectors. Triple mutants tR>A, tR>L, tR>K were commercially synthesized (GenScript) and cloned into MIEG.

Production of infectious retroviral supernatant were carried out using polyethylenimine to transfect 5 μg of retroviral plasmids into Plat-E packaging cells. Supernatant was collected at 48h and 72h after transfection. Retroviral supernatant containing various C/EBP constructs was added to dKO-B cells together with 8 ug/mL polybrene (Sigma). Cells were infected by spinoculation at 2000 rpm at 37°C for 1 hour, following by overnight incubation. Cells were then washed and resuspended in fresh medium. Transdifferentiation of dKO-B cells was executed as described (Cirovic *et al*., 2017).

### Cell transfection

HEKT-293 cells were transfected with C/EBPα WT and mutant expression vectors in the absence or presence of hWDR-HA or hBrm-HA (SMARCA4) as indicated, using Polyethylenimine according to the manufacturer’s instructions (PEI, Polysciences #24765-2).

### Flow cytometric analysis and sorting

After harvesting at indicated time points, cells were washed in cold FACs buffer (2% FCS, 2 mM EDTA in PBS) and incubated with Fc-blocking reagent (1:200, TruStain FcX, anti-mouse CD16/32, BioLegend). After washing, cells were proceeded to labelling with fluorescence-conjugated antibodies at 4°C for 30 minutes in the dark. Stained cells were washed and resuspended in FACs buffer containing propidium iodide (PI, BD Biosciences) for live-dead discrimination. Samples were measured using LSRFortessa analyzer. Cells were sorted using FACSAria II/III instruments (BD Biosciences). Gentamicin was added to buffer and post-sort cell culture medium at final concentration 10 μg /mL. Sorted cells were spun down in gentle cycle (700 rpm, 7 minutes) and resuspended in fresh medium. Measurement was recorded using FACSDiva and analyzed using Flowjo^TM^ software v8.

Antibodies used in this study include:

transdifferentiation panel CD115 APC (eBioscience, #17-1152-80), CD11b PE (BD Pharmingen, #557397), CD19 PE-Cy7 (eBioscience, #25-0193-81), Ly6G BV421 (BioLegend, #127628);

bone marrow sorting panel CD117 (cKit) PE-Cy7 (eBioscience, #25-1171-82), Streptavidin PECy-7 (BioLegend, #405206), Ly6A/E (Sca-1) PE (BD Pharmingen, # 553336), APC-conjugated Lineage cocktail containing CD3e (BD Pharmingen *#*553066), Ly6C/Ly6G (BD Pharmingen #553129), CD11b (BD Pharmingen #553312), Ter119 (BioLegend #116211), B220 (BD Pharmingen # 553092), CD11c (BD Pharmingen #550261), CD5 (eBioscience #17-0051-82), CD115 (BioLegend #135509);

colony characterization panel CD16/32 PE (BD Pharmingen #561727), CD117 (cKit) PE-Cy7 (eBioscience #25-1171-82), CD34 AlexaFluor 647 (BioLegend #152205), Ly-6G AlexaFluor 700 (BioLegend #127621), Ly-6C APC-Cy7 (BioLegend #128025), F4/80 Pacific Blue (BioLegend #123123), CD115 Brilliant Violet 605 (BioLegend #135517), CD11b Brilliant Violet 711 (BioLegend #101241), CD11c PerCP-Cy5.5 (BioLegend, # 117327).

### Isolation of bone marrow cells and LSK cells

Bone marrow-derived cells were isolated from femur, tibia and part of hip joints of 8–12-week-old Cebpa^fl/fl^Cebpb^fl/fl^ mice. Bones were flushed with cold PBS using a syringe and a 24-gauge needle (for serial replating of total cKit-enriched bone marrow cells) or crushed in cold PBS (for serial replating of LSK and GMP cells) under sterile condition. Cell suspension was filtered through a 70 μM cell strainer before incubation with red blood cell lysis solution (Miltenyi) for 8-10 minutes on ice. Cells were washed and cultured in cytokine supplemented-IMDM or resuspended in PBS for further processing.

### Enrichment of cKit^+^ cells

Isolated bone marrow cells were incubated with biotin-conjugated cKit antibody (BioLegend, #105803) for 20 minutes at 4°C. After washing, anti-biotin magnetic beads (Miltenyi Biotec, #130-090-485) were added to cell suspension (20 μL beads per 10^7^ cells) and incubated for 30 minutes at 4°C. Magnetically labelled cells were washed and resuspended in max. 10^9^ cells/mL and passed through equilibrated selecting columns (LS Column, Miltenyi) mounted on a MidiMACs or a QuadroMACs separator. Cell-loaded columns were washed 3 times with MACs buffer, transferred to a 15 mL centrifuge tube and eluted with 1 mL MACs buffer. Cells were then washed and cultured in cytokine supplemented-IMDM or prepared for further processing.

### Colony serial replating assay

Colony serial replating assay was performed on C/EBPα p30 (WT or mutants) infected, GFP^+^ sorted bone marrow cells. Directly after sorting, cells were resuspended in cytokine supplemented-IMDM at 5 x 10^4^ cells/mL. Cells mixture was diluted in Methocult medium (MC3434, STEMCELL Technology) to obtain a final concentration of 5000 cells/mL. Semi-solid MC3434 cell mixtures were plated at 1 mL mixture/well in a 6-well meniscus-free dish (SmartDishTM, STEMCELL Technology) using a 3cc syringe and a blunt-end 16-gauge needle (STEMCELL Technology). Colony dishes were scanned using EVOS^TM^ FL^TM^ auto imaging system (Thermo Fisher Scientific) every 7 days for 4 passages (day 7, day 14, day 21 and day 28) at 4x magnification. Colonies were counted and classified manually based on morphology.

To replate, cell suspension in three replicates were pooled, spun and washed with PBS. Colonies were well-suspended to obtain single cells suspension and diluted to 5 x 10^4^ cells/mL in cytokine supplemented-IMDM. Cell suspension was added to MC3434 aliquots and seeded as described above. Leftover cells of day 7 plates (Plate 1) were subjected to further analysis, including flow cytometric analysis, proliferation analysis and cytospin.

### Proliferation assays

Growth curves were determined by cell counting and WST-1 assay. For cell counting, 10^5^ cells were seeded in 1mL cytokine supplemented IMDM into a 24-well plate in triplication. Cells were count every 24 hours by staining with Trypan blue and using a Neubauer chamber.

For WST-1 assay, 10^4^ cells were seeded in 100 μL cytokine supplemented IMDM into a 96-well plate in triplication. Every 24 hours, 10 μL of WST-1 reagent (Roche) was added to each well (1:10 dilution) and incubated at 37°C. After 60 minutes, absorbance of 450 nm wavelength was measured using an iMark microplate absorbance reader (Bio-Rad). A blank control wells, which contained equivalent cell culture medium was used for normalization of OD values. Proliferation rate was determined by dye-dilution assay using CellTrace Violet (ThermoFisher Scientific). Cells were processed following manufacturer’s suggested procedure. An aliquot of the culture suspension was drawn at indicated time points and labelled with antibodies including Ly6C, Ly6G, CD115 and CD11b. Labelled cells were measured as described; PI was used for live-dead determination.

### May-Grünwald/Giemsa staining

Cells were spun onto glass slides using an Aerospray Slide Stainer cytocentrifuge (Wescor) at 500 rpm for 5 minutes. Air-dried slides were fixed with methanol before immersed into May-Grünwald solution for 5 minutes. After washing, slides were immersed in freshly prepared Giemsa solution for 35-45 minutes. Stained slides were rinsed, air-dried, and mounted with mounting solution (Roti-Histokitt II). Stained cells were observed under the microscope and captured at multicolor mode (EVOS, Invitrogen).

### Protein extraction, immunoprecipitation and western blot

Production of whole cell lysates and immunoprecipitation of WT or mutant C/EBPα proteins were performed as previously described (Kowenz-Leutz *et al*., 2010). Briefly, cells were lyzed (20 mM Hepes pH 7.8, 150 mM NaCl, 1 mM EDTA pH 8, 10 mM MgCl2, 0,1% Triton X-100, 10%Glycerol, protease inhibitor cocktail (Merck), 1 mM DTT, 1 mM PEFA bloc (Böhringer), and immunoprecipitation was performed with antibodies as indicated for 2 h at 4^0^C. Immunoprecipitated proteins were collected on Protein-G Dynabeads (Invitrogen #10004D), separated by SDS-PAGE (Mini PROTEAN TGX, 4-15%, Bio-Rad #5671084) and immunoblots were incubated with antibodies (HA, Covance #MMS-101R; Flag, Sigma #F3165), as indicated and visualized by ECL (GE Healthcare, UK).

### WDR5 interaction mapping on spot-synthesized tiling p30 C/EBPα peptides

To map post-translational dependent WDR5 interaction with the three arginine residues R140, R147, R154 contained in the p30 C/EBPα N-terminus a protein interaction screen on a custom PepSpot cellulose membrane (PRISMA; JPT Peptid Technologies, Berlin) (Dittmar *et al*., 2019) was performed using tiling peptides of 13 aa length spanning the critical Arginine residues (see also Supplemental Figure S4).

Arginine residues within peptides were modified to Rme2as, Rme2sym, citrulline or alanine. Binding specificity was controlled by histone H3 or Mlll1 peptides, as indicated. A bacterially expressed and purified human WDR5-N-His construct (aa24-334) was incubated with the membrane, as described (Dittmar *et al*., 2019; Ramberger *et al*., 2021). The membrane was blocked for 20 min with Rotiblock (Roth #A151.1) and incubated with an anti-WDR5 specific antibody (Abcam #ab56919) in Rotiblock for 20 min at RT. The membrane was washed twice for 5 min with TBST 0.01% Tween and incubated with anti-mouse HRP antibody (Invitrogen #31431) for 15 min at RT. After washing twice with TBST 0.01% Tween for 5 min the filter was developed with ECL (Amersham) and signals determined and quantified on a Licor scanner (C-DiGit Blot Scanner).

### Bulk RNA-sequencing

After virus infection (as in **Figure 1**), EGFP^+^ cells were sorted at day 4 p.i. and subjected to bulk RNA-sequencing. Cell pellets were resuspended in RNA lysis buffer (RNeasy Kit, Qiagen) and store in -80°C condition. Samples were in biological quadruplicates, all samples harvested in different batches were processed altogether using RNAeasy Kit (Qiagen). For RNA-seq, 1 μg of total RNA was used. The concentration of extracted total RNA was measured using Qubit 3 Fluorometer (Thermo Fisher Scientific). Quality and integrity of RNA were measured using the Eukaryote Total RNA Nano assay on Bioanalyzer 2100 (Agilent Technologies). Preparation of barcoded mRNA-seq library, sequencing using NextSeq500 platform (Illumina) with paired-end reading at 75 bps read-length were performed at the EMBL Genomics Core Facility (Germany).

### Bioinformatic analysis of mRNA sequencing data

Raw sequencing reads were aligned to the mm10 reference genome using STAR version 2.7.9a. Subsequently, htseq-count version 0.1 was applied for gene expression quantification. Normalization and differential gene expression were performed with DESeq2 using an adjusted p-value cutoff of 0.05 and an absolute fold change of 2 as parameters defining differential expression. Functional analysis was done with gProfiler using GO:BP and KEGG as an annotation resource.

### Statistical analysis

Data were analyzed for statistical significance by tests indicated at each figure using GraphPad Prism 10.1.1.

## ACKNOWLEDGMENTS

We thank Hans-Peter Rahn and Kirstin Rautenberg (MDC, Berlin, Germany) for the flow cytometry support; Alexandra Patmanidi (MDC, Berlin, Germany) and Vladimír Beneš (Genomics Core Facility, EMBL, Heidelberg, Germany) for the RNA-seq support.

## AUTHOR CONTRIBUTIONS

Nguyen, L.T.: Investigation, Formal analysis, Original Draft, Visualization, Validation; Zimmermann, K.: Investigation, Formal analysis, Data curation, Validation; Kowenz-Leutz, E.: Investigation, Formal analysis, Methodology, Resources, Validation, Supervision; Dörr, D.: Formal analysis, Methodology; Schütz, A.: Investigation, Formal analysis; Schönheit, J.: Investigation, Formal analysis; Mildner, A.: Investigation, Methodology, Supervision; Leutz, A.: Conceptualization, Supervision, Project administration, Funding acquisition, Methodology, Investigation, Review & editing.

## DECLARATION OF INTERESTS

The authors declare no competing interests.

## Notes

### Competing Interest Statement

The authors have declared no competing interest.

