## Supplementary Figures for "Arginine methylation of the p30 C/EBPα oncoprotein regulates progenitor proliferation and myeloid differentiation"

**Figure S1 (related to Figure 1)**

**A.** Gating strategy of flow cytometric analyses. Debris and doublets were excluded using FSC and SSC gating, live construct-expressing cells ($PI^{neg}EGFP^{+}$) were targeted for marker analysis. Successfully transdifferentiated cells are $CD19^{dim/neg}CD11b^{+}$, percentages of which were used for graphs in **Figure 1C, 1D** and **Figure S1B, S1C**.

**B.** Percentage of $CD19^{dim/neg}CD11b^{+}$ cells. Data are shown as mean ± SEM. Significance was determined by Turkey's test (mixed-effects analysis with Geisser-Greenhouse corrections), multiple comparisons against p42 C/EBPα are shown, n.s > 0.05, *p ≤ 0.05, **p ≤ 0.01, ***p ≤ 0.005, ****p ≤ 0.001. Note that cells expressing p42 C/EBPα were growth arrested and gradually disappeared from the culture.

**C.** Percentage of $PI^{neg}EGFP^{+}$ cells. Data are shown as mean ± SEM. Significance was determined by Turkey's test (mixed-effects analysis with Geisser-Greenhouse corrections), multiple comparisons against p42 C/EBPα are shown. Note that from day 0 to day 4, percentage of MIEG and p30 C/EBPα expressing cells remained approximately similar, while cell counts of p42 C/EBPα cells dropped drastically (observation, data day 0-4 not shown).

**Figure S2 (related to Figure2)**

**A.** Gene ontology (GO) enrichment analysis of differentially expressed genes in comparison to vector control. Top enriched GO-Biological processes (GO-BP) were shown in order of increasing -log10(p-value). For a completed list of enrichment terms, see Supplementary Table 1. Abbreviations: pos. (positive), neg. (negative), reg. (regulation), nucl. (nuclear), cat. (catabolic), proc. (process), inflam. (inflammatory), inv. (involved).

**B.** Heatmap presenting pair-wise differential gene expression analysis. Genes with adjusted p-value 0.05, |FC|>2 in at least one comparison are listed.

**Figure S3 (related to Figure 3)**

**A.** Colony formation during four passages of replating total c-Kit-enriched bone marrow cells. Colony numbers identified by counting of image scans. Colony formation rates per $10^4$ cells from three technical replicates; one representative experiment is shown. All biological independent experiment yielded similar results.

**B.** Gating strategy of cell sorting to enrich for hematopoietic stem cells, **LSK** ($Lin^{neg}cKit^{+}Sca\text{-}1^{+}$) and granulocyte monocyte progenitors **GMP** ($Lin^{neg}cKit^{+}Sca\text{-}1^{neg}FcgR^{+}$).

**C.** Growth curve by colorimetric assay (WST-1) using cells from Plate #1 of total c-Kit enriched bone marrow (as shown in Figure 3A, S3A). Data are shown as mean ± SEM, significance was determined by two-way ANOVA analysis followed by Dunnett's multiple comparisons test, only significance between WT and other constructs is shown by asterisks in matching colors, *p ≤ 0.05, **p ≤ 0.01, ***p ≤ 0.005, ****p ≤ 0.001. Results of one experiment with three technical replicates are shown. Three independent experiments showed similar results.

**D.** Dye dilution assay using cells from Plate #1 of LSK replating assay. Fluorescence intensity of $EGFP^{+}$ CellTrace-Violet-stained cells was measured at indicated time points using flow cytometric analysis. Tinted bars mark unstained population, numbers indicate percentage of cells with complete dilution of fluorescence dye. Histogram shows modal counts in each sample.

**Figure S4 (related to Figure 4)**

**A.** Giemsa-May-Grunewald staining of cytospin cells from pooled Plate #1 colonies. Note the distribution of neutrophil lineage cells (polymorphic nuclei) in comparison to monocyte lineage cells (full, rounded nucleus). Scale bar 50 µm.

**B.** Gating strategy to analyze cell type distribution on colony Plate #1 using flow cytometry. Colonies from three technical replicates were pooled, an aliquot of $10^6$ cells were subjected to FACs analysis. Standard gating was similar to **Figure 1**, $EGFP^+$ cells were further gated based on Ly6C expression. $Ly6C^+$ cells were further stratified into neutrophils ($Ly6G^+$) or monocytes ($CD115^+$). Percentage of each population, as shown in red, was plotted in **Figure 4C, 4D**.

**C.** t-SNE plots of FACs analysis in **Figure 4C, 4D**. Heatmap represented expression level of Ly6C, with $Ly6C^-$ cells indicated. $Ly6G^+$ and $CD115^+$ population were also shown. For t-SNE plotting, 5000 events of each sample were randomly subtracted using DownSample plugin of Flowjo. Further sample-concatenation and dimensional reduction was applied using t-SNE tool (Flowjo). Gating was determined based on unstained and fluorescence-minus-one controls. Representative data from one experiment are shown.

**D.** Dye dilution assay, as shown in **Figure S3B**, with Ly6C staining integrated. Intensity of CellTrace-Violet dye was determined separately in $Ly6C^-$ and $Ly6C^+$ population. Red tinted line indicated maximum division, black tinted line indicated minimum division.

**E.** C/EBPα sequence (aa 132–175) and tiling peptide sequences underneath. Each peptide was spot synthesized as WT or in modified configuration (Rme2as, Rme2sym, Cit, A), as indicated below.

**F.** Interaction of purified WDR5 with the PepSpot cellulose membrane according to the PRISMA protocol. Briefly, the PepSpot blot with synthesized C/EBPα peptides was incubated with 10 ug/ml recombinant N-His-WDR5. After washing, the PepSpot blot was incubated with anti-WDR5 and subsequently with anti-mouse horseradish peroxidase antibody and visualized by ECL.

**G.** Coomassie stained SDS-gel of recombinant N-His-WDR5 aa 24-334. Lane 1 (crtl): extract of uninduced cells, lane 2 (ind): IPTG induced cells, lane 3 (purif): Ni-agarose purified WDR5.

**H.** Quantification of WDR5 interactions (from blot as shown in F.) using Licor scanning. Quantitative data from peptides 16-20, 21-25, and control peptides (peptide # 46-51; H3 and KMT2A) are shown as a bar graph.

### Supplement Figure 1

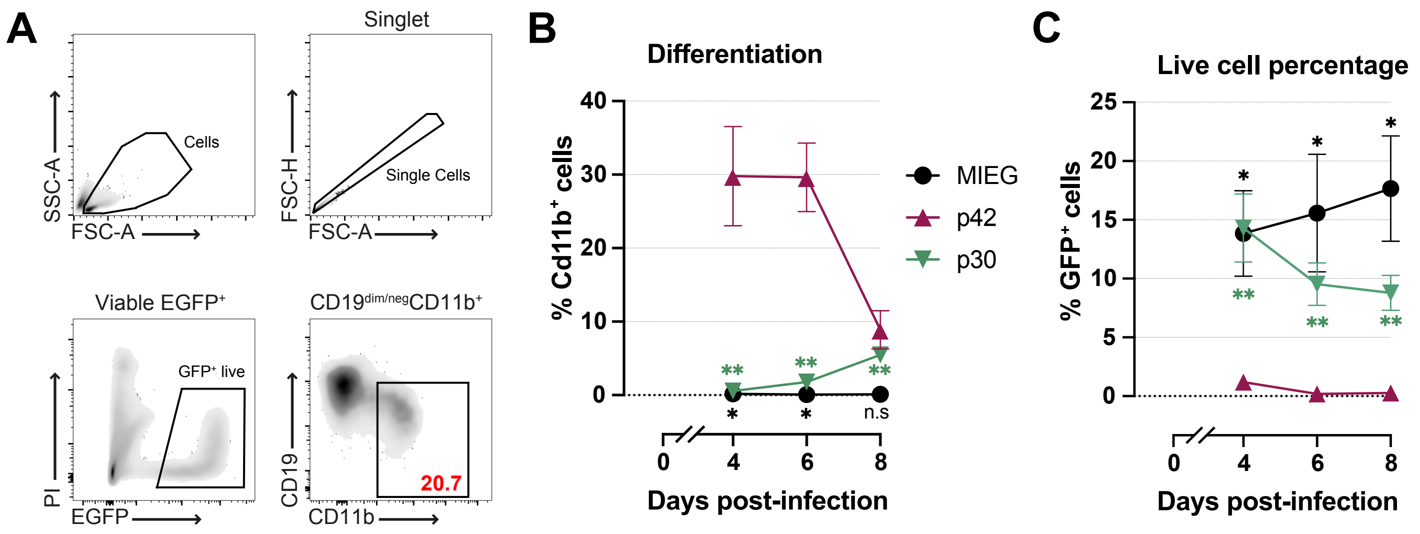

### Supplement Figure 2

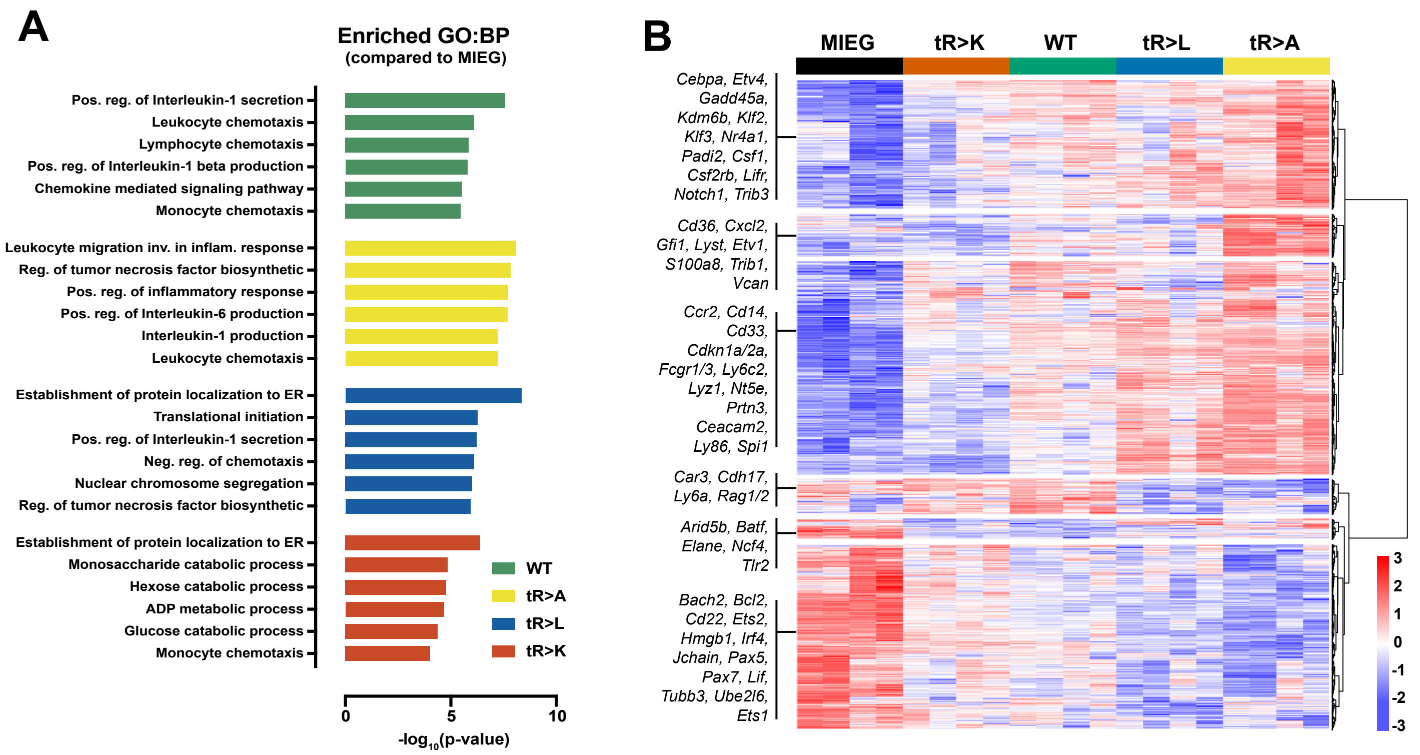

### Supplement Figure 3

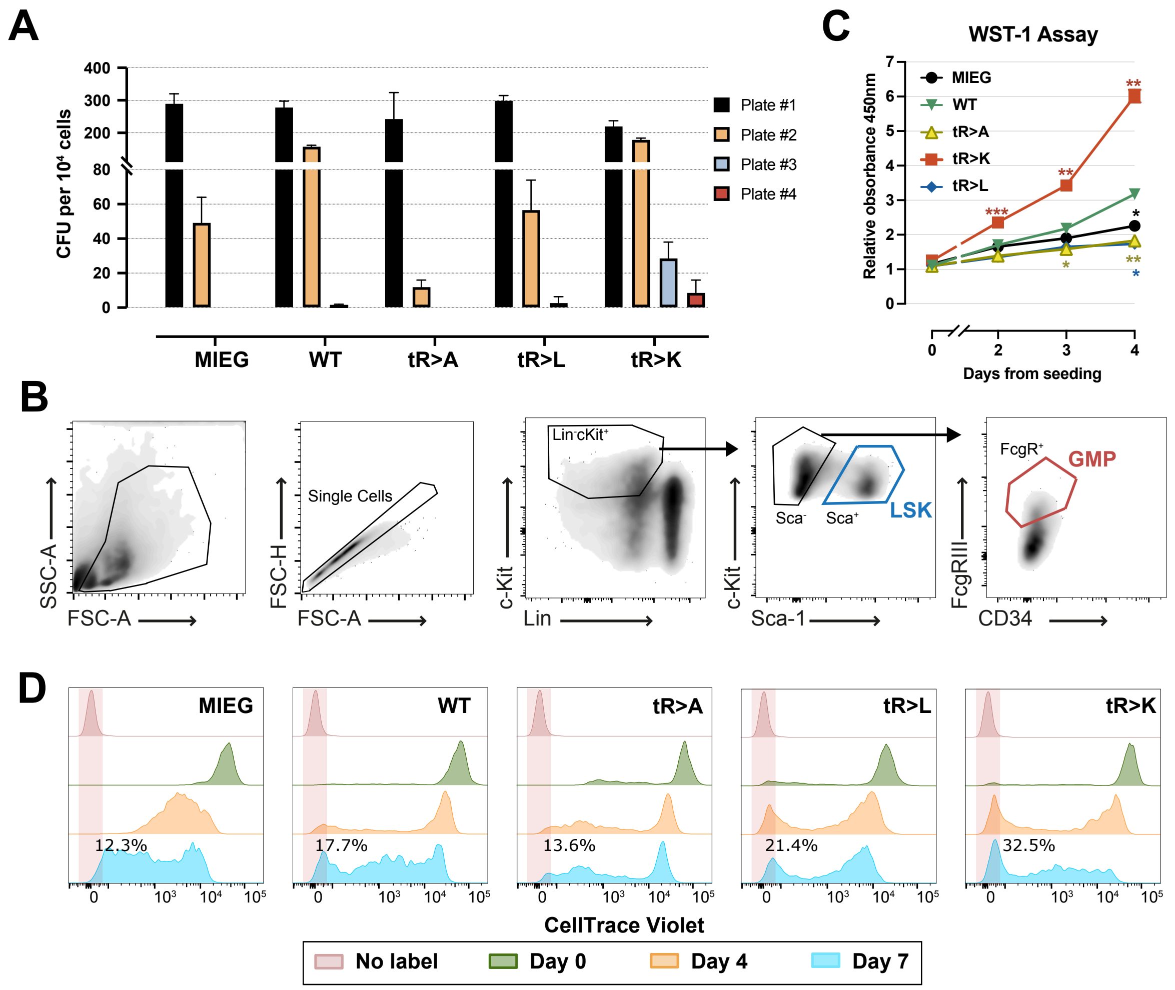

### Supplement Figure 4

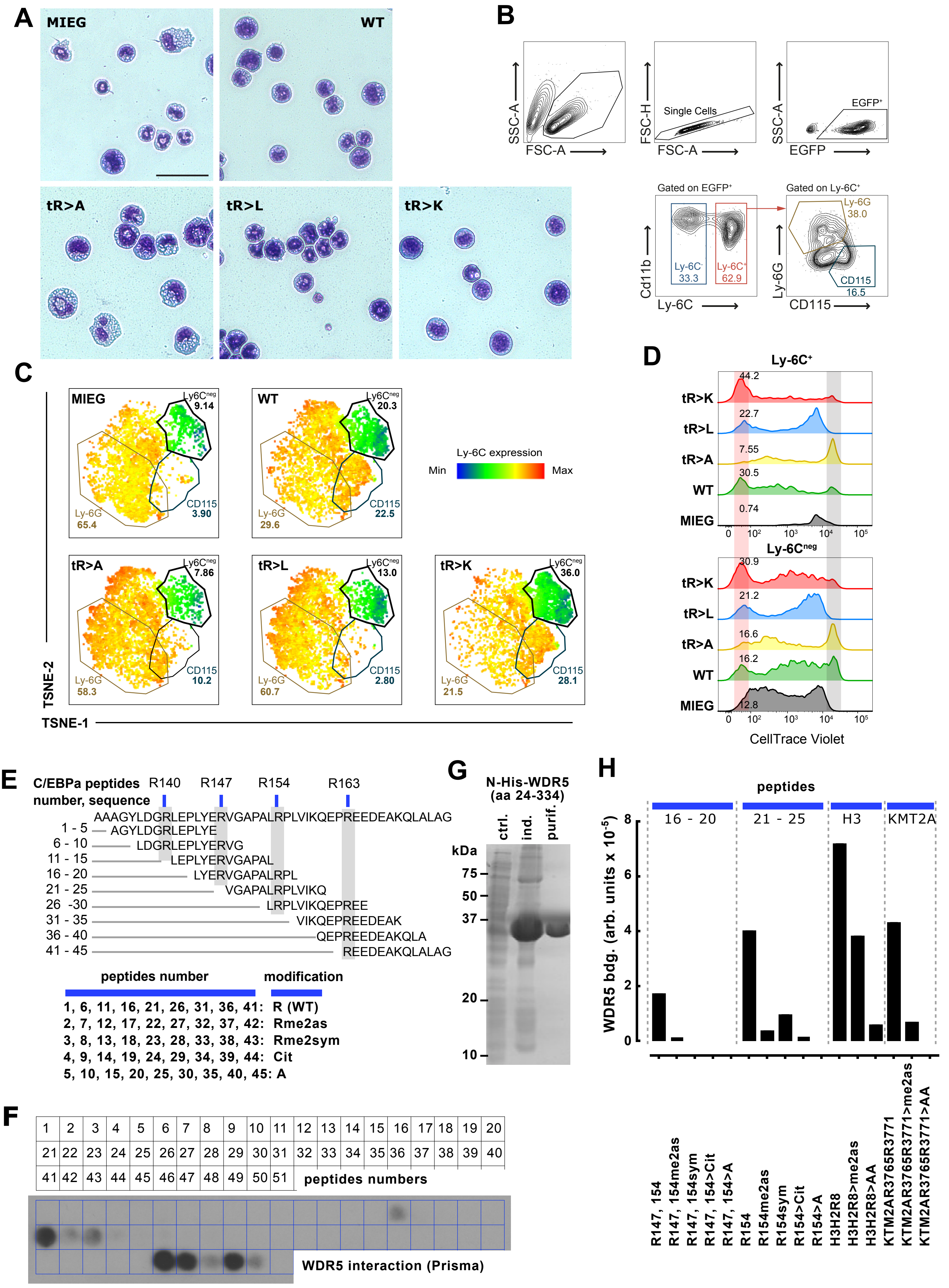
